# Divergent warning patterns influence male and female mating behaviours in a tropical butterfly

**DOI:** 10.1101/2023.08.05.552099

**Authors:** Chi-Yun Kuo, Lina Melo-Flóres, Andrea Aragón, Morgan M. Oberweiser, W. Owen McMillan, Carolina Pardo-Diaz, Camilo Salazar, Richard M. Merrill

**Affiliations:** Division of Evolutionary Biology, Ludwig-Maximilians-Universität München, Grosshaderner Strasse 2, 82152 Planegg-Martinsried, Germany; Smithsonian Tropical Research Institute, Gamboa 0843-03092, Panamá; Department of Biological Science and Environmental Biology, Kaohsiung Medical University, 100 Shih-Chuan 1 Rd, 80708 Kaohsiung City, Taiwan; Department of Biology, Faculty of Natural Sciences, Universidad de Rosario, Carrera 24 # 63C-69, Bogotá, 111221, Colombia

**Keywords:** assortative mating, computer vision, mate choice, mate preference, reproductive isolation

## Abstract

Traits under divergent ecological selection that also function during mating can be important in maintaining species boundaries. Few studies have considered mutual mate choice, where both males and females base mating decisions on the same trait. We investigated the contribution of visual preference to assortative mating between an aposematic butterfly *Heliconius cydno* and its sympatric relative *H. melpomene*. Wing colouration in *Heliconius* butterflies evolved as a warning signal but also functions as a mating cue. We predicted that both sexes of *H. cydno* contributed to assortative mating by exhibiting visual preference towards conspecific wing colouration. We analysed published and new data from preference experiments, in which males were presented with conspecific and *H. melpomene* females. We also recorded female responses and mating outcomes in choice experiments, involving conspecific males with either the original white or artificially painted red forewing bands. Both sexes of *H. cydno* responded more positively towards the conspecific colouration, yet male colouration did not predict mating outcomes in female choice experiments. As courtships are initiated by males in butterflies, our findings suggest that female visual preference might be of secondary importance in preventing gene flow between *H. cydno* and *H. melpomene*.

## Introduction

Divergent natural selection is a major force driving speciation. It is increasingly recognized that traits under divergent ecological selection can also contribute to non-random mating (also known as the ‘magic traits’ or ‘multiple effect’ traits [1,2]). This mechanism is important as it offers an evolutionary solution to the conundrum of maintaining associations between traits under divergent ecological selection and those that cause assortative mating. These magic traits can be used by either sex as the basis for mate choice, but the contribution of such traits to reproductive isolation depends on the mating system and might be different between sexes [3,4] (but see [5]).

In ecologically divergent species where both sexes can exert mate choice, examining the role of mate preference in only one sex might misinform the behavioural underpinnings of reproductive isolation in two ways. First, the strength and direction of male and female preference can differ within species (e.g., [6]). Second, the existence of mate preference does not guarantee that such preference would contribute to reproductive isolation. For example, in *Gasterosteus* threespine sticklebacks, ecologically divergent benthic and limnetic species differ in body size and colouration. Females of each ecotype use body size and to a lesser extent colour as mating cues, leading to assortative mating [7]. Interestingly, even though males have been observed to court differentially, it does not predict mating outcome and is unlikely to contribute to reproductive isolation [7].

For most systems, however, we have only limited knowledge on whether and how male and female mate preference may act in synergy to prevent interspecific gene flow (Table S1). This is true in *Heliconius* butterflies, a radiation characterized by rapid convergent and divergent evolution in wing colour patterns. This group of butterflies are classic examples of Müllerian mimicry, where multiple chemically defended species evolve the same warning colour patterns to signal their unprofitability toward predators [8]. Closely related *Heliconius* species often have distinct colour patterns, which are maintained by strong selection against hybrids [9,10]. Colour pattern differences also constitute an important premating barrier as males almost invariably prefer females with the same colour pattern [11–13]. However, although studies have suggested female choice may contribute to assortative mating in *Heliconius* butterflies, the magnitude of female preference to conspecific male colour patterns has rarely been quantified (e.g., [6,14]).

Here, we test for both male and female visual preferences in two *Heliconius* butterflies. *Heliconius cydno* and *H. melpomene* coexist in neotropical America, have distinct colour patterns from mimicking different *Heliconius* species [10](Figure S1). The two species are closely related but mate assortatively, based at least in part on male preferences for conspecific colour pattern [15,16]. We first quantified male visual preference in *H. melpomene* and *H. cydno* using both previously published and new data. To test whether female visual preference may also contribute to reproductive isolation, we performed choice experiments in which *H. cydno* females encountered conspecific males that differed in wing colouration.

## Materials and Methods

All statistical analyses were performed in R (version 4.2.2 [17]).

### Male visual preference experiments

#### Experimental setup

Mature virgin males (> 5 days old) of *H. cydno* and *H. melpomene* were housed together in a cage (2 × 4 × 2 m) and were provided daily with flowers of *Psychotria, Psiguria* and *Lantana* plants, as well as sugar water with dissolved pollen as food source. We gave males individual markings by painting dots on the dorsal side of the forewings with a black marker (Copic, Tokyo, Japan); each male had a unique combination of dots (e.g., one dot on the right forewing and two dots on the left forewing). Dot combinations were reused when existing males were replaced by new individuals. To quantify visual preference, we simultaneously presented the male butterflies with one *H. cydno* and one *H. melpomene* model made of freshly dissected wings in 4-hour daily trials and filmed their behaviour with mounted GoPro cameras (Hero 5, San Mateo, CA, USA). Cage and camera setup can be found in the Supplementary Methods. We used custom scripts to screen for the occurrence of potential preference behaviours from video recordings by identifying specific flight patterns as described in [18].

#### Statistical analyses

In addition to data from this study, we also re-analysed data from [11] and [16]. [11] recorded male visual preference as the number of male courtships towards *H. melpomene* and *H. cydno* model butterflies in groups of males without differentiating between individuals. They also used models with real dissected wings and those with paper wings. For data from [16] and the present study, the number of preference behaviours towards each female was recorded for individuals. [16] used live females. We used generalized linear (mixed) models (GL(M)M) to test whether male *H. cydno* and *H. melpomene* displayed more courtships or preference behaviours towards females of their own colour patterns. In all datasets, we used the numbers of courtships towards each model as binomial response variables. For data from [11], we performed binomial GLM with species and model type (i.e., real wings vs. paper wings) as fixed-effect predictors, as including group as a random factor resulted in singular-fit model with trivial amount of variance. For data from [16] and the present study, we tested the existence of male visual preference using GLMM, with species as a fixed-effect predictor and individual as a random factor.

### Female visual preference experiments

#### Experimental setup

We painted the wings the day before an experiment to minimize the effect of paint on males’ flight ability and behavior, even though the paint normally dried within minutes after application. Each male was then placed individually in a cage (2 × 4 × 2 m) and was allowed to acclimate overnight. On the morning of the experiment, the experimental female was placed in a popup cage and put in the cage for ten minutes to acclimatize before being released. We recorded female responses as either acceptance or rejection based on [14] (Table S2).

#### Behavioural trials

In order to test how forewing colouration in male *H. cydno* might affect female responses and mate preference, we first conducted “no-choice” experiments, in which a one-or two-day old virgin female was paired with a conspecific virgin male aged at least 10 days. We used females of this specific age because this is the age at which mating typically takes place in the wild, when females are most receptive [19]. Each male was randomly assigned to either the “red” or “white” treatment prior to the experiments. Males assigned to the red treatment had their white forewing bands coloured with a red marker (Copic Ciao R27, Tokyo, Japan) on both the dorsal and ventral sides, whereas males in the white treatment had their forewing bands painted in the same way but with a colourless blender marker (Copic Ciao 0, Copic, Tokyo, Japan). The choice of marker colour was determined by multispectral image analyses developed by [20] in ImageJ [21] in order to match the red colour of *H. melpomene* (see Supplementary Methods). The colourless and the red markers were identical in their chemical content, differing only in colour.

Female responses to male courtships were recorded continuously for up to 2 hours or until mating occurred. We recorded each female response as either acceptance or rejection following [14](Table S2). We then monitored whether mating occurred between a pair every 30 minutes from 8:00 am to 5:30 - 6:00 pm for two days [22]. Each individual was only used once to ensure independence of trials. We performed the experiments described above at the Smithsonian Tropical Research Institute in Panama during January to April of 2019 and at the Estación Experimental José Celestino Mutis of Universidad del Rosario in Colombia during February to April of 2020.

In a second experiment, we performed “choice” experiments in Colombia, in which a virgin *H. cydno* female aged 1-3 days was presented with two virgin males aged at least 7 days. The age difference between the two males was at most two days. We manipulated the colourations of the two males so that one male had the white forewing bands and the other the red forewing bands. We checked whether mating occurred and the colour of the mated male every 30 mins from 8:00 am to 6:00 pm for two days. We did not record the interactions between the sexes in these experiments.

#### Statistical analyses

We also used GLMM to test whether female *H. cydno* responded more positively towards control males, and whether female responses determined mating outcomes. Whenever appropriate, we calculated standardized effect sizes (odds ratio or Cohen’s D) and the associated 95% confidence interval when there was at least moderate evidence for a fixed effect. We interpreted the magnitude of Cohen’s D following [23]. Additional details of the statistical procedures are available in the Supplementary Methods.

## Results

We used the language of evidence when reporting results [24].

There was very strong evidence from all datasets that males of both species were more likely to perform courtships and preference behaviours towards models of their own colour patterns, with odds ratios larger than 20 (Table 1, Figure 1). Similarly, we found very strong evidence that females reacted to colour pattern differences. Although females largely exhibited rejection behaviours towards approaching males (Figure 2A), the proportion of acceptance behaviours was higher towards “white” males (P < 0.0001 z = 4.41, Table 2, Figure 2A), with a medium effect size (Cohen’s D = 0.45). However, there was no evidence that female response, male courtship, or male colouration predicted the occurrence of mating in these no-choice trials (P > 0.34, Table 2, Figure 2B). Indeed, in 10 out of the 21 two-choice experiments, the female mated with the red male. A sign test revealed no evidence that male colouration was associated with mating outcome (probability of success = 0.476 [95% confidence interval: 0.257 – 0.702], P-value = 1).

**Table 1.**
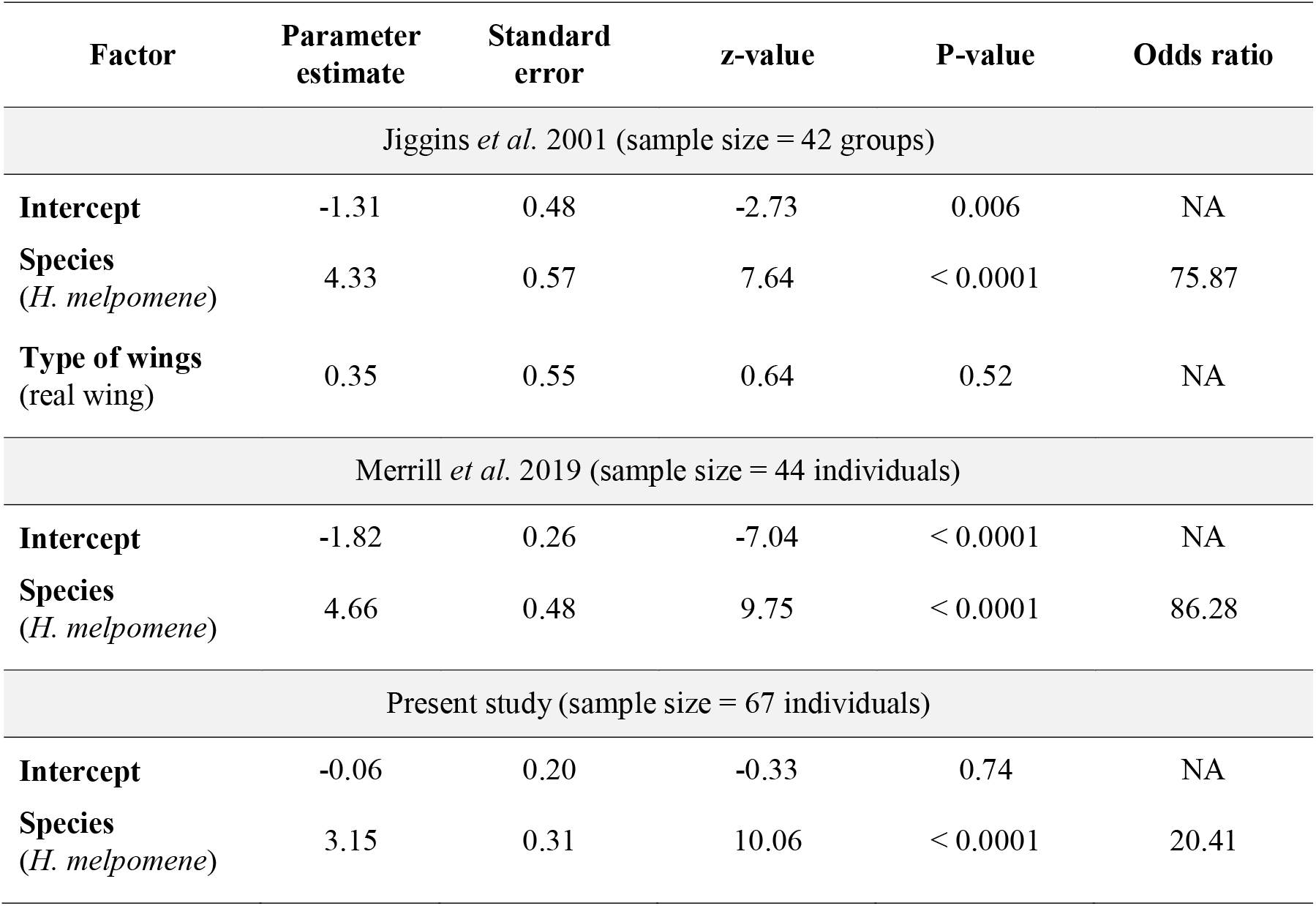
Statistical results of male visual preference from the three datasets.

| Factor | Parameter estimate | Standard error | z-value | P-value | Odds ratio |
| --- | --- | --- | --- | --- | --- |
| Jiggins *et al.* 2001 (sample size = 42 groups) |  |  |  |  |  |
| **Intercept** | -1.31 | 0.48 | -2.73 | 0.006 | NA |
| **Species**<br>(*H. melpomene*) | 4.33 | 0.57 | 7.64 | < 0.0001 | 75.87 |
| **Type of wings**<br>(real wing) | 0.35 | 0.55 | 0.64 | 0.52 | NA |
| Merrill *et al.* 2019 (sample size = 44 individuals) |  |  |  |  |  |
| **Intercept** | -1.82 | 0.26 | -7.04 | < 0.0001 | NA |
| **Species**<br>(*H. melpomene*) | 4.66 | 0.48 | 9.75 | < 0.0001 | 86.28 |
| Present study (sample size = 67 individuals) |  |  |  |  |  |
| **Intercept** | -0.06 | 0.20 | -0.33 | 0.74 | NA |
| **Species**<br>(*H. melpomene*) | 3.15 | 0.31 | 10.06 | < 0.0001 | 20.41 |

**Table 2.** Statistical results examining the variation in female response and mating outcome in no-choice experiments.

| Factor | Parameter estimate | Standard error | z-value | P-value | Cohen's D |
| --- | --- | --- | --- | --- | --- |
| Proportion of female acceptance behaviour |  |  |  |  |  |
| **Intercept** | -1.03 | 0.74 | -1.39 | 0.16 | NA |
| **Male colour**<br>(white) | 0.82 | 0.19 | 4.41 | < 0.0001 | 0.45 |
| Mating outcome |  |  |  |  |  |
| **Intercept** | -0.62 | 0.68 | -0.90 | 0.37 | NA |
| **Species**<br>(*H. cydno chioneus*) | 0.33 | 0.86 | 0.38 | 0.70 | NA |
| **Male colour**<br>(white) | 0.29 | 0.74 | 0.40 | 0.69 | NA |
| **No. of female acceptance behaviours** | 0.05 | 0.06 | 0.75 | 0.46 | NA |
| **No. of male courtships** | -0.03 | 0.03 | -0.96 | 0.34 | NA |

**Figure 1.**
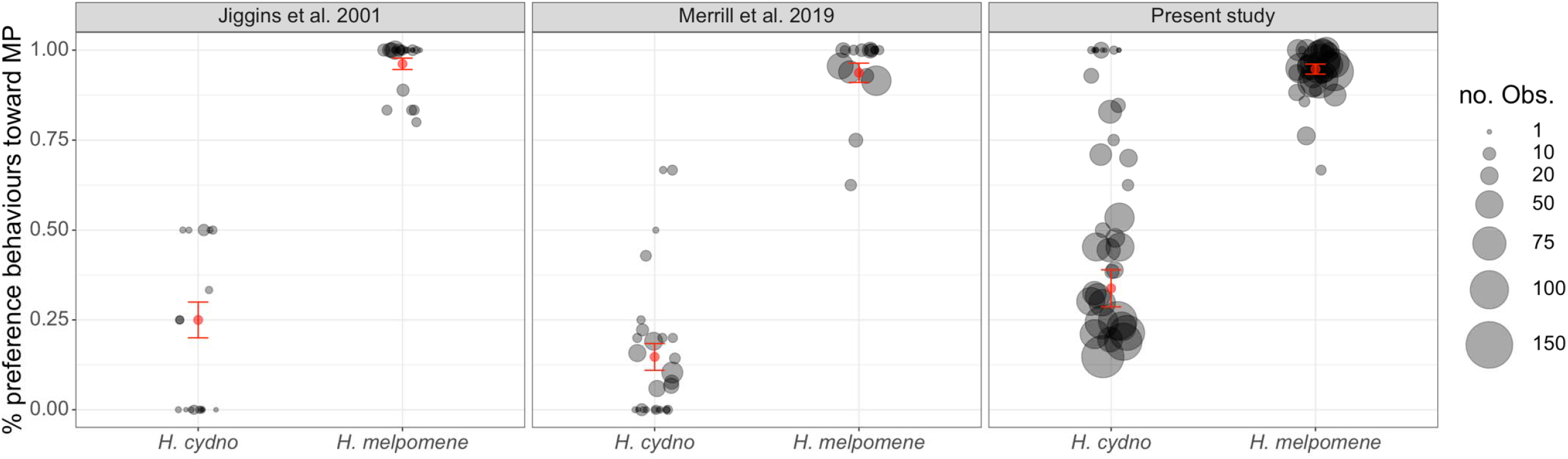
Male visual preference of *H. melpomene* and *H. cydno* from three datasets, quantified as proportion of preference behaviours towards the female *H. melpomene* (MP) model. Each circle represented a group of males in [11] and an individual male in the other two datasets. Size of the circle reflects the number of observations per group or individual. Red dots and error bars denote means and standard errors.

**Figure 2.**
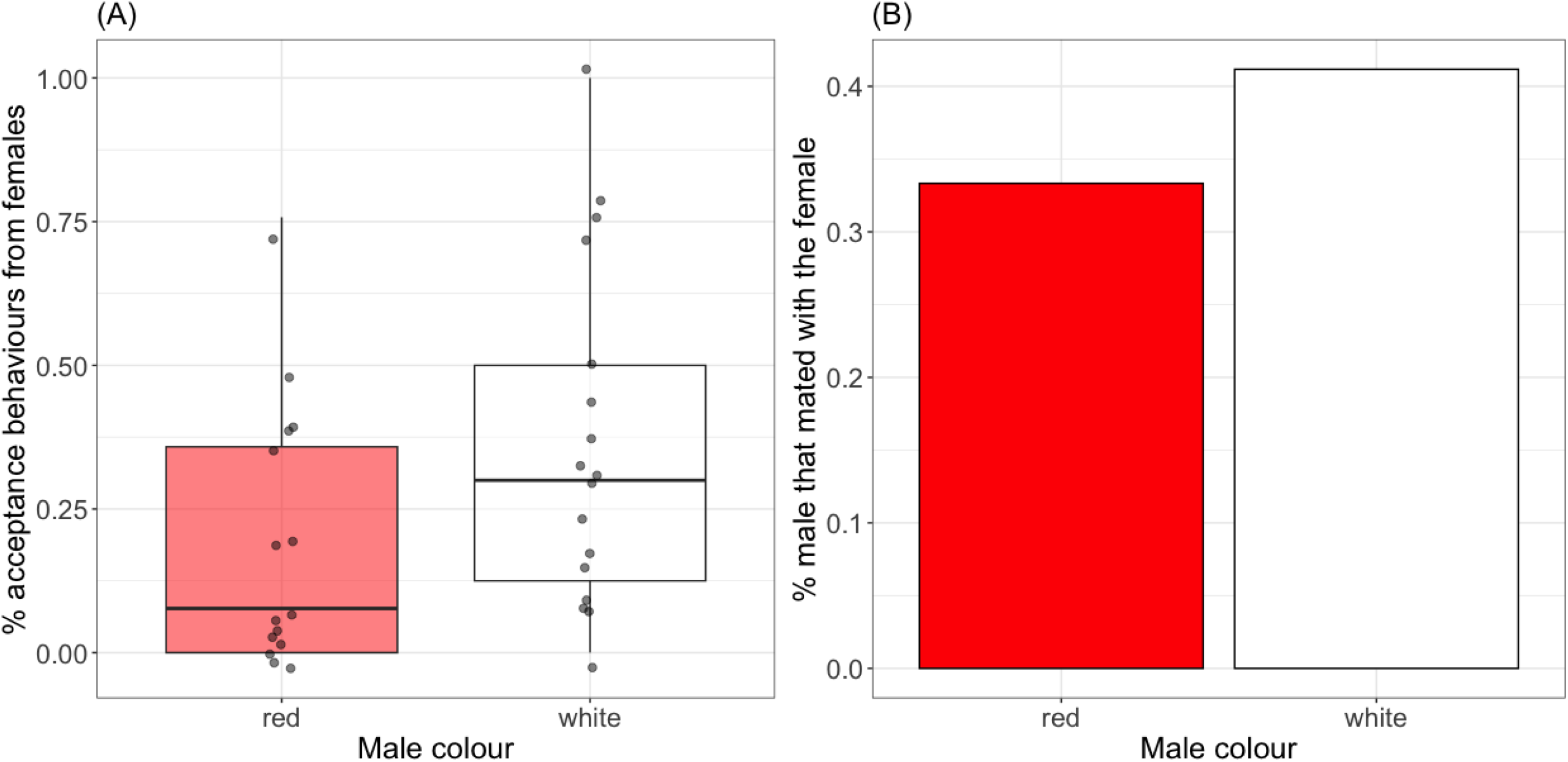
(A) Proportion of female acceptance behaviour in each male colour treatment. Females responded more positively towards males from the white group. (B) Proportion of males that mated with the female in each wing colour treatment. The difference in male wing colours did not affect the likelihood of mating. Results were based on 15 males from the red group and 17 males from the white group.

## Discussions

### Wing colouration influences mating behaviours of both sexes in H. cydno

Both sexes of *H. cydno* responded more positively towards conspecific wing colouration. Males on average showed more interests towards *H. cydno* models when a *H. melpomene* model was also present. Even though most responses towards male courtships were rejections, females did display more acceptance behaviours towards males with the original wing colouration. These findings show that both sexes base their mating behaviours on wing colouration, which is an honest warning signal under strong disruptive selection between sympatric *Heliconius* species. The role of wing colouration in mating preference has been extensively studied for male *Heliconius* (e.g., [6,11,18,22]). In females, the importance of visual information on female mating behaviours has only been examined in *H. numata*, in which females prefer wing patterns that are dissimilar to their own [6], and in sympatric *H. erato* and *H. himera*, in which females did not use visual cues for mate choice [25]. A previous study reported that female choice acted as a reproductive barrier in *H. cydno*, but their experimental design does not differentiate the contribution of wing colouration versus information from other sensory modalities (e.g., chemical signals)[14]. Our results demonstrated the relevance of male wing colouration in eliciting differential female responses in *H. cydno*.

### Male wing colouration did not determine mating outcomes in H. cydno

Despite the influence of male wing colouration on female response, it did not appear to determine mating outcomes in *H. cydno*. Interestingly, neither more courtship effort nor more female acceptance behaviours predicted the occurrence of mating in this study. A plausible explanation might be that mating outcome is determined by nonvisual signals or cues, such as the compositions of male sex pheromone, which are known to affect female choice in *H. melpomene, H. erato, H. himera*, and *H. timareta* [25,26]. In addition, individual variation in male sex pheromone compositions has been reported in *Heliconius* butterflies [27]. It is possible that individual males in our experiments differed intrinsically in their pheromonal profiles, which led to variation in perceived suitability by the females and, ultimately, to differential mating outcomes. Testing this possibility could be a fruitful next step.

### Implications on reproductive isolation between sympatric H. melpomene and H. cydno

Courtship in butterflies is initiated by males, so that male preference acts before female preference. Earlier acting barriers are expected to have a stronger effect on overall reproductive isolation, as they may limit gene flow before other barriers can act [28]. Nevertheless, female preference could still contribute to assortative mating when male preference towards conspecific females is not absolute. In *H. cydno*, males direct on average only 70-85 percent of their courtships towards conspecific females. However, our findings showed that even though female *H. cydno* did respond more positively towards males with white forewing bands, male colouration did not play a pivotal role in determining mating outcomes. This implies that female visual preference of *H. cydno* might not be important in contributing to prezygotic isolation with *H. melpomene*. This does not refute the role of female choice in maintaining the species boundaries between these two species, as females could still use other sensory information for mate choice (e.g., [29,30]). We acknowledge that our experiments did not directly test female visual preference when given a choice between *H. melpomene* and *H. cydno* colourations. However, given the fact that female preference can only be investigated as responses to male courtships and that visual preference is very strong in *H. melpomene* males, such experiment may not be practical to perform.

## Data accessibility

Data and R code associated with this manuscript are available at https://github.com/thekuolab/cydno_female_preference

## Authors’ contributions

C.-Y. K.: conceptualization, data collection, formal analysis, investigation, methodology, visualization, writing – original draft, review and editing; L. M.-F, A. A., and M. M. O.: data collection, methodology; W. O. M.: funding acquisition, investigation; C. P.-D. and C. S.: investigation; R. M. M.: conceptualization, formal analysis, funding acquisition, investigation, methodology, writing – original draft, review and editing

All authors gave final approval for submission and agreed to be held accountable for the work performed therein.

## Acknowledgements

We would like to thank Oscar Paneso, Cruz Batista, and Isabel Leon for help with butterfly care. We are grateful to *Ministerio de Ambiente*, Panamá (permit no. SE/AP-14-18) and *Autoridad Nacional de Licencias Ambientales*, Colombia (permit no. 530) for permissions to collect butterflies.

