## Supplemental Materials for "Divergent warning patterns influence male and female mating behaviours in a tropical butterfly"

**I. Supplementary Methods**

**1. Setup for recording and screening for male preference behaviours**

There were six camera stations evenly spaced in the experimental cage that also allowed us to mount a model butterfly about 70 cm above the ground. Models were made of *H. cydno* and *H. melpomene* females sacrificed immediately after eclosion and pinned with wings fully open. We washed the wings of model butterflies with hexane to remove potential pheromone residuals prior to the experiments. To minimize the chance of males becoming desensitized towards model females, we randomly assigned the hourly location of the models such that each model was presented at four different stations in a 4-hour trial. was mounted 50 cm above each model to capture motions of butterflies approaching the model. The pairing between cameras and models was also randomized daily. Raw videos were recorded in 1080p pixel resolution, a frame rate of 60 per second, a shutter speed of 1/480 seconds, 1600 ISO, and a Narrow field of view (focal length 28 mm) with the model at the center.

The main script extracted all butterfly movements from raw recordings and rendered each incident of movement into a five-second footage. The program then concatenated all movements captured from one camera into a single video file. This method allowed us to collect high-throughput behavioural data from a large number of males on a daily basis (four hours per camera per day). We then examined the condensed videos and recorded a male as exhibiting associative behaviour towards a model if it exhibited either an “approach” or a “courtship” behaviour or when the male made physical contact with the model [1].

**2. Multispectral image analyses for marker choice**

To select the red colour that best approximates *H. melpomene* under butterfly vision, we purchased eight different red markers (Copic Ciao R05, R14, R17, R27, R29, R35, R46 and RV29, Tokyo, Japan) and performed multispectral image analyses developed by [2] in ImageJ [3] to determine which marker most resembled the colour of the red forewing band in *H m. rosina*. We first coloured eight distinct white areas on a *H. cydno* wing with each of the marker. We then took photographs of the coloured *H. cydno* wing and a *H. melpomene rosina* forewing with a Nikon Nikkor D7000 camera (Nikon, Melville NY, USA) with a visible light (range allowed) or a UV (range allowed) filter (company, city). A 40% grey standard (company, city) was included in each photograph for colour calibration. The visible light and UV photos of each wing were combined to generate a multispectral image, which the software then converted to cone catch models based on the visual sensitivity and relative abundance of cone receptors for species in the *H. melpomene* clade [4,5]. Based on the cone catch model results, we calculated pairwise just noticeable differences (JND) between the red forewing band in *H. melpomene rosina* and each of the eight coloured areas on the *H. cydno* wing, using a Weber fraction of 0.05. A JND value less than 2 is conventionally considered as indistinguishable by the visual system [2]. The marker R27 (Cadmium Red) had the lowest pairwise JND (1.78) and was therefore the marker we used to alter the forewing colours in experimental *H. cydno* males.

**3. Additional information on statistical analyses**

For female visual preference, we first tested whether male coloration had an effect on female responses. To do so, we used the number of acceptance and rejection responses as binomial response variable, male colour treatment as a fixed-effect factor and subspecies (*H. cydno chioneus* vs. *H. cydno cydno*) as a random-effect factor in generalized linear mixed models (GLMM). Variance inflation factor scores did not reveal any severe collinearity among predictors (all values < 1.03). Male courting effort, measured as the total number of courtships, did not differ between colour treatments (Kruskal-Wallis test: 𝜒^2^ = 0.25, df = 1, P = 0.62) and was therefore not included in the statistical model. We then tested whether species, male treatment, and the number of receptive female responses were associated with mating outcome of each trial by constructing a logistic GLM with mating outcome as the binary response variable. There was no significant collinearity among predictors (variance inflation factor values < 1.3). Visual examination of the data did not reveal any obvious sign of interactions between predictors. We examined whether male colorations had an effect on the likelihood of mating from the two-male experiments with a sign test, using data from the 21 out of the 41 trials in which mating occurred.

All statistical models were validated with residual diagnostics [6,7] using the DHARMa package [8] before interpretation.

**II. Supplementary Tables and Figure**

**Table S1.** Examples of traits that are both the targets of divergent ecological selection and mating cues for assortative mating based on [9], with added information on the contributions of male and female preference to reproductive isolation. We only included examples from [9] in which mate preference was explicitly tested for at least one sex.

| **System** | **Divergent forms** | **Putative magic trait(s)** | **Tests of male and female preference as causes of reproductive isolation** | **Reference** |
| --- | --- | --- | --- | --- |
| *Gasterosteus*  freshwater sticklebacks | Limnetic and benthic forms | Body size | Male (tested, negative)  Female (tested, positive) | [10] |
| *Gambusia* fishes | Predator and predator-free forms | Body shape | Male (not tested)  Female (tested, positive) | [11] |
| *Heliconius* butterflies | Different mimetic forms | Colour pattern | Male (tested, positive)  Female (not tested) | [12–14] |
| *Dendrobates pumilio*  Poison-dart frogs | Different aposematic forms | Colour and colour pattern | Male (not tested),  Female (tested, positive) | [15,16] |
| *Geospiza fortis*  Darwin’s finches | Ecologically divergent individuals within a population | Beak morphology | Male (not tested)  Female (tested, positive) | [17] |
| *Lycaeides* butterflies^a^ | Species in different habitats | Wing colour pattern | Male (tested, positive)  Female (not tested) | [18] |
| *Loxia curvirostra* crossbill birds | Different “call” types | Foraging rate | Male (not tested)  Female (tested, positive) | [19] |
| *Mormyridae* electric fish | Different electric discharges | Electric organ discharge | Male (not tested)  Female (tested, positive) | [20] |

^a^ The cause of divergent selection was not identified.

**Table S2.** Description and categorization of behaviours recorded between male and female *H. cydno,* following [21]*.*

| **Sex** | **Behaviour** | **Description** | **Category** |
| --- | --- | --- | --- |
| Males | Chase | Male follows female closely while both are flying | Courtship |
|  | Hover | Male hovers over perched female | Courtship |
|  | Mating attempt | Male lands next to female and bends abdomen towards hers | Courtship |
| Females | Close wings | Perched female holds wings closed. | Acceptance |
|  | Open wings | Perched female opens wings and holds them there. | Rejection |
|  | Flutter | Perched female rapidly opens and closes wings while lifting abdomen and, usually, exposing abdominal scent glands. | Rejection |
|  | Fly | Perched female flies away from male (including taking off from a perched position | Rejection |

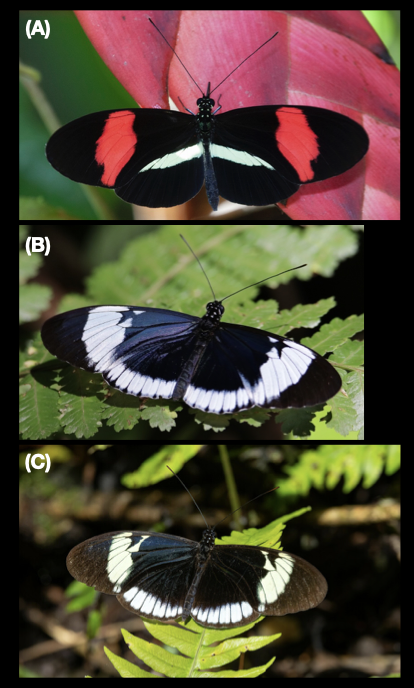

**Figure S1.** *Heliconius melpomene* *rosina* (A) and *H. cydno* used in this study. We used *H. cydno chioneus* in Panama (B) and *H. cydno cydno* in Colombia (C). Images are from iNaturalist (https://www.inaturalist.org).
